# Investigating the *cis-*Regulatory Basis of C_3_ and C_4_ Photosynthesis in Grasses at Single-Cell Resolution

**DOI:** 10.1101/2024.01.05.574340

**Authors:** John Pablo Mendieta, Xiaoyu Tu, Daiquan Jiang, Haidong Yan, Xuan Zhang, Alexandre P. Marand, Silin Zhong, Robert J. Schmitz

## Abstract

While considerable knowledge exists about the enzymes pivotal for C_4_ photosynthesis, much less is known about the *cis-*regulation important for specifying their expression in distinct cell types. Here, we use single-cell-indexed ATAC-seq to identify cell-type-specific accessible chromatin regions (ACRs) associated with C_4_ enzymes for five different grass species. This study spans four C_4_ species, covering three distinct photosynthetic subtypes: *Zea mays* and *Sorghum bicolor* (NADP-ME), *Panicum miliaceum* (NAD-ME), *Urochloa fusca* (PEPCK), along with the C_3_ outgroup *Oryza sativa*. We studied the *cis-*regulatory landscape of enzymes essential across all C_4_ species and those unique to C_4_ subtypes, measuring cell-type-specific biases for C_4_ enzymes using chromatin accessibility data. Integrating these data with phylogenetics revealed diverse co-option of gene family members between species, showcasing the various paths of C_4_ evolution. Besides promoter proximal ACRs, we found that, on average, C_4_ genes have two to three distal cell-type-specific ACRs, highlighting the complexity and divergent nature of C_4_ evolution. Examining the evolutionary history of these cell-type-specific ACRs revealed a spectrum of conserved and novel ACRs, even among closely related species, indicating ongoing evolution of *cis*-regulation at these C_4_ loci. This study illuminates the dynamic and complex nature of CRE evolution in C_4_ photosynthesis, particularly highlighting the intricate *cis-*regulatory evolution of key loci. Our findings offer a valuable resource for future investigations, potentially aiding in the optimization of C_3_ crop performance under changing climatic conditions.

## Introduction

Photosynthesis is one of the most critical chemical reactions on the planet whereby CO_2_ is metabolized into glucose. Plants have evolved numerous variations of photosynthesis. The most common type of photosynthesis uses the enzyme ribulose 1,5-biphosphate carboxylase oxygenase (RuBisCO) which combines CO_2_ with a five carbon compound ribulose 1,5-biphosphate to create 3-phosphoglyceric acid. This three-carbon compound is then used in a redox reaction within the Calvin Benson cycle, where sucrose is made. The production of this three-carbon compound is what gives this type of photosynthesis, C_3_, its name. However, although widely evolved and found in many crop plants, C_3_ photosynthesis struggles to perform in hot, arid conditions. In non-ideal conditions, O2 can competitively bind the RuBisCO active site, causing the formation of a toxic intermediate, and reducing photosynthetic efficiency and plant performance (1). Due to increasing temperature caused by anthropogenic climate change, this reduction in photosynthetic capacity for key crop plants poses a major agricultural challenge (2). However, other types of photosynthesis have evolved in hotter conditions and offer a model to potentially alter key C_3_ crop plants to be more efficient.

The C_4_ photosynthetic pathway is an example of a modified style of photosynthesis that is able to perform in hot conditions. In brief, C_4_ typically works by sequestering key photosynthetic enzymes into two different compartments in the leaf made up of different cell types. These two cell types/compartments are bundle sheath (BS) cells, which in C_4_ plants generally form a concentric ring around the vasculature, and mesophyll (MS) cells, which make up large portions of the non-vascularized leaf internal cells (3). In the MS, CO_2_ is imported, and converted to bicarbonate (HCO3-) by the enzyme carbonic anhydrase (CA). Bicarbonate is then converted to a four-carbon molecule oxaloacetate (OAA) by the O2-insensitive phosphoenolpyruvate carboxylase (PEPC). This OAA molecule made of a four-carbon compound (where C_4_ derives its name) is finally converted into a stable metabolite, either malate or aspartate. This intermediate molecule is then transported to the BS where it undergoes a decarboxylation process, by one of three different types of decarboxylases, NAD-dependent malic enzyme (NAD-ME), NADP-dependent malic enzyme (NADP-ME), or phosphoenolpyruvate carboxykinase (PEPCK). This decarboxylation reaction releases a CO_2_ molecule that enters into the Calvin Benson cycle. The generation and processing of intermediate molecules in cellular compartments allows for concentrated levels of CO_2_ to interact with RuBisCO, reducing the inefficiencies mentioned above. Additional types of C_4_ photosynthesis have been observed which don’t rely on division of metabolites between MS and BS cell-types, but instead rely on using dimorphic chloroplast instead as in the species *Bienertia sinuspersici* (4,5). Current C_4_ crops such as maize (*Zea mays*), sorghum (*Sorghum bicolor*), pearl millet (*Cenchrus americanus)*, foxtail millet (*Setaria italica*), and broomcorn millet (*Panicum miliaceum)* excel in their ability to operate in adverse conditions.

Although the evolution of C_4_ photosynthesis is a complex process, there is tantalizing evidence that engineering C_3_ crops to do C_4_ photosynthesis might be possible. One piece of evidence that points to this is that C_4_ photosynthesis has evolved independently 65 times in different lineages of plants (6). These results indicate that most plant lineages have the genetic material capable of evolving into C_4_ photosynthesizers. The *Poaceae* lineage of grasses exemplifies this, as C_4_ photosynthesis has evolved independently at least 18 times (7). Interestingly, all of these species use the same core C_4_ enzymes and steps, but many use different decarboxylation enzymes as mentioned above (8–10). Furthering this hypothesis is the fact that many C_4_ related genes originally evolved from either C_3_ photosynthetic genes or key enzymes critical in core metabolism (11,12). For instance, PEPC is a key metabolism enzyme in the glycolytic pathways of the Krebs Cycle, with some copies being important in guard cell metabolism (13–15). Instead of novel gene content being the main driver of C_4_ photosynthesis, it’s more likely due to the correct timing and compartmentalization of key enzymes into specific cell types (16–18). This raises the question, how is gene expression of these key C_4_ enzymes regulated? Moreover, as C_4_ has evolved multiple times convergently, have similar regulatory networks and paradigms been co-opted to alter when and where these key genes are expressed?

*Cis-*regulatory elements (CREs) are key players in gene regulation, as they both fine tune expression and provide cell-type specificity (19–22). In brief, these regions operate as binding sites for transcription factors (TFs). Transcription factors are proteins which are able to alter transcription by binding DNA sequences and recruiting transcriptional machinery which can either increase or decrease transcription (23). Thus TFs are able to significantly change molecular phenotypes. Previous work has shown that CREs could be key players in the transition to C_4_ photosynthesis. This was demonstrated by taking C_4_ genes from *Z. mays* and transforming them into *Oryza sativa,* a C_3_ species (24,25), which revealed that CREs from *Z. mays* genes were able to drive cell-type-specific expression in MS in *O. sativa* (24,25).

Additional analyses have implicated CREs as drivers in the evolution of C_4_ photosynthesis. In the genus of plants *Flaveria,* which contains both C_4_ and C_3_ plants, one key difference in C_4_ plants was a specific CRE driving gene expression in MS cells. This 41 bp motif named *Mesophyll expression module 1* is critical for cell-type-specific expression of *PEPC* in MS cells, a critical first step in the C_4_ pathway (19,26). Finally, four conserved non-coding sequences were identified to be critical in MS-specific expression of *PEPC* in monocots (27). Furthermore, a recent cross-species study examining the binding sites of GLK, a conserved TF regulating photosynthetic genes, revealed that CREs can undergo rapid changes and result in diverse gene expression patterns without the need of altering the TF itself (28). These findings show that CREs are important genetic elements that plants use for the evolution of C_4_ photosynthesis.

Although some CREs critical for cell-type-specific expression of key photosynthetic genes have been identified, they’ve been restricted to those nearby the transcriptional start sites. This is due, in part, to the challenge of identifying CREs genome wide, as well as limitations in the isolation of BS and MS cells which is labor intensive and challenging. However, a recent study used a multi-omic approach in *Z. mays* BS and MS cells and found CREs genome-wide that might be critical in the cell-type-specific regulation of genes (29). One example is the identification of a potential distal CRE ∼40 kb upstream of *SULFATE TRANSPORTER4* (*ZmSFP4)*, a BS-specific sulfate transporter (29). These results highlight the complexity of identifying loci involved in *cis* regulation. Identifying all CREs associated with C_4_ loci is critical in enhancing our understanding of *cis* regulation of key C_4_ genes, and would greatly enhance attempts at engineering C_3_ crops. During the evolution of C_4_ photosynthesis, it’s unclear whether these CREs have been pre-established during evolution and co-opted for C_4_ photosynthesis or if they evolved independently numerous times. Understanding the ways in which *cis* regulation evolves to control timing and cell-type-specific expression of C_4_ photosynthesis genes would greatly assist efforts in engineering C_3_ plants to be more C_4_ like.

To investigate the role of CREs and their potential contribution in controlling key C_4_ genes, we used single-cell indexed Assay for Transposase Accessible Chromatin sequencing (sciATAC-seq) to identify cell-type-specific CREs from five grass species representing diverse C_4_ subtypes, as well as an additional C_3_ outgroup. We investigated the cell-type specificity of both the core C_4_ enzymes, and those which are unique to each photosynthetic subtype. Further, we identify CREs of C_4_ genes, and find previously unknown cell-type-specific CREs that might be critical in C_4_ gene expression. We find that some of these regulatory regions appear not just conserved in a single C_4_ subtype, but in all of the C_4_ species we studied. Finally, we leverage these data to find transcription factor binding motifs enriched in MS and BS cell types and use these motifs to catalog these regulatory loci.

## Results

### Identification and Annotation of Cell Types in Diverse Species

To investigate CREs in BS and MS cells potentially important in C_4_ photosynthesis, we generated replicated sciATAC-seq libraries for four different C_4_ species, comprising three different C_4_ subtypes NADP-ME (*Z. mays, S. bicolor*), NAD-ME (*Panicum miliaceum)*, and PEPCK (*Urochloa fusca),* and a C_3_ outgroup species (*O. sativa*) (**Figure 1A**). Libraries were filtered for high-quality cells by first pseudo-bulking the sciATAC-seq libraries, and identifying accessible chromatin regions (ACRs). Using these ACRs, per nuclei quality metrics were then calculated such as fraction of reads in peaks, transcriptional start site enrichment, and total integration events per nucleus (**Methods**). Nuclei found to have a high proportion of organellar reads were also removed, with values being adjusted on a per library basis (**Methods**). Clustering of cells was done on genomic bins, and with additional cells removed that had a high correlation with *in-silico* generated doublets, and clusters were removed that were skewed towards one replicate by greater than 75% (**Methods**). After filtering on per nucleus quality metrics, we identified 16,060 nuclei in *Z. mays*, 15,301 nuclei in *S. bicolor*, 7,081 nuclei in *P. miliaceum*, 19,110 nuclei in *U. fusca,* and 5,952 nuclei in *O. sativa* (**Supplemental Figure 1, Supplemental Table 1**).

**Figure 1:**
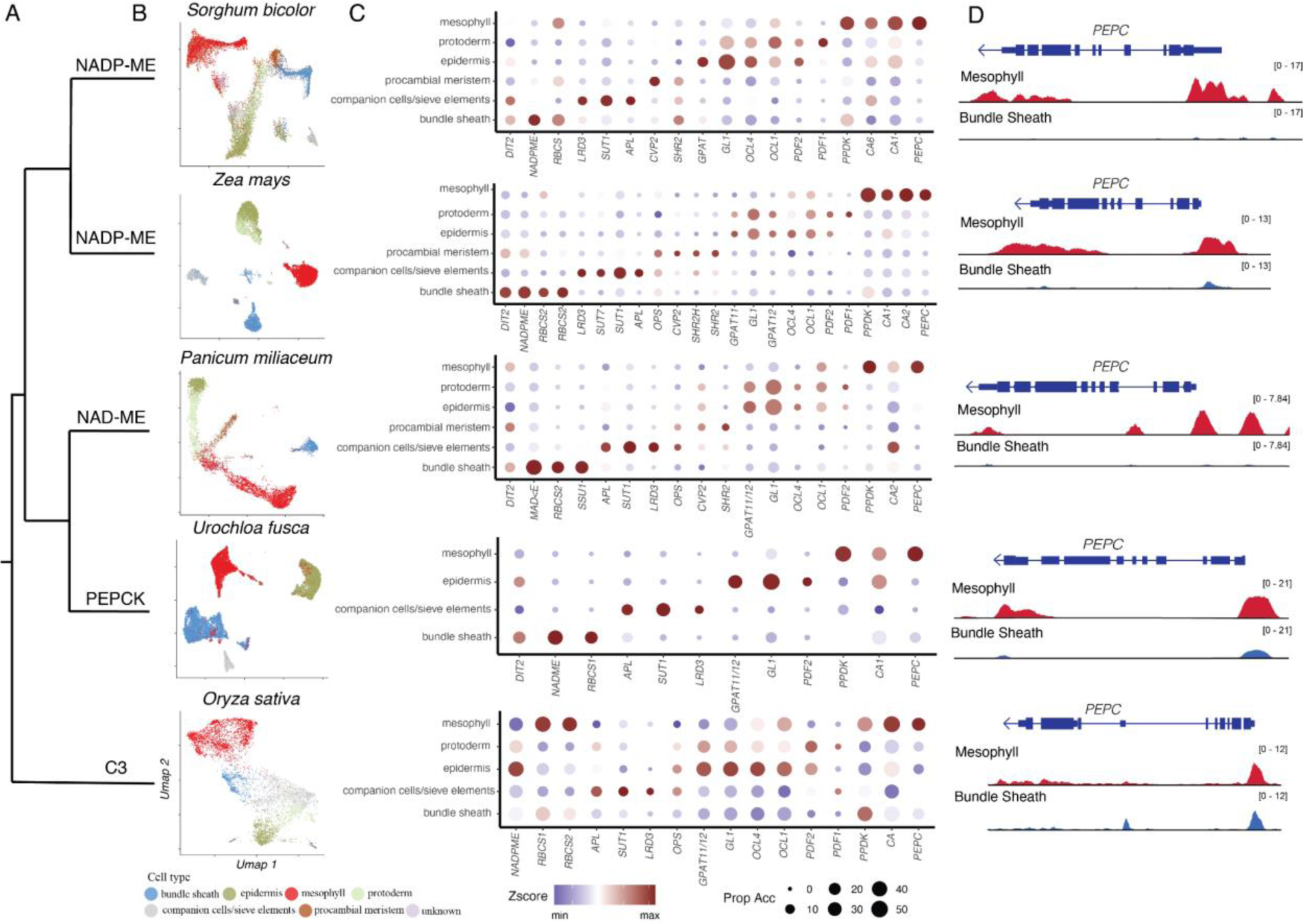
Annotation of cell types in diverse grass species at single-cell resolution **A)** A phylogeny indicating the relationship of various C_3_ and C_4_ photosynthesizers sampled. In this sample, two NADP-ME subtypes are represented, one NAD-ME subtype, a PEPCK subtype, as well as a C_3_ species. **B)** UMAP embedding showing the annotation for each species. A cell type legend is below. **C)** Dotplots for various marker genes used to annotate each species. The y-axis represents cell types, and the x-axis is a list marker genes used to annotate different cell types. The size of each circle is proportional to the number of cells within that cell type that showed chromatin accessibility of the marker. Color is z-score transformed values across clusters of gene chromatin accessibility across the clusters. **D)** Screenshots of the *PEPC* locus for all sampled species. For each screenshot, the top track shows the protein coding, the red track is chromatin accessibility of MS cells, and the blue track is the chromatin accessibility of the BS cells.

Due to variation in genome size and content, cell-type annotation for each dataset was done independently using the reference genome for each species (**Figure 1B**). We used multiple approaches to annotate cell types. Orthologs of key marker genes from *Z. mays* and *O. sativa* were identified using a phylogenetics based approach (**Methods**). This allowed for the identification of marker genes for specific cell types in a cross species context. To gauge gene activity of these marker genes, gene body chromatin accessibility was used as a proxy for expression (**Figure 1D**) (21,30). Cell-type annotation was done manually taking into consideration marker gene chromatin accessibility, marker enrichment in clusters, as well as ontological relationships between cell types (**Supplemental Figure 2-19**). Due to the lack of marker genes for many cell types in plants, as well as the challenge of annotating a broad sample of species, we reduced resolution of our annotation across our datasets to ensure accurate comparisons between variable species (**Figure 1B**).

Deeper exploration of the list of marker genes from *Z. mays* showed conservation of gene body chromatin accessibility in markers for certain cell types (**Supplemental Table 2-3**). As expected, for the C_4_ plants, *RIBULOSE BISPHOSPHATE CARBOXYLASE SMALL SUBUNIT1 (RBCS1) and RIBULOSE BISPHOSPHATE CARBOXYLASE SMALL SUBUNIT2* (*RBCS2*) were enriched in BS cells compared to MS cells (**Figure 1C**), a pattern that was not found in *O. sativa.* Additionally, *PEPC1* showed MS-specific chromatin accessibility in all of the C_4_ species sampled (**Figure 1D**). Additionally, we found conservation of marker genes like *SUCROSE TRANSPORTER 1* (*SUT1*) in companion cells and sieve elements, and *GLOSSY1* (*GL1*) in epidermis cells, indicating that these historically described marker genes are likely important in this diverse set of species. This analysis provides a first examination of core-C_4_ marker genes’ chromatin accessibility across a diverse sample of plant species at cell-type resolution.

### Chromatin Accessibility of Core C_4_ Enzymes Shows Similar Cell-Type Bias, but Differing Evolutionary Origins

We measured the chromatin accessibility bias of the C_4_-associated enzymes. Due to the diverse nature of the plants sampled, and the C_4_ photosynthetic subtypes, we separated enzymes into core- and subtype-specific groups. This list comprised nine core C_4_ enzymes, and nine variable enzymes. These enzymes were assigned to one of these two groups based on if they are found in all C_4_ subtypes (core) or are specific to only one or two subtypes (variable). One example of a core enzyme is carbonic anhydrase, which is used to generate bicarbonate from CO_2_, as well as for the regeneration of phosphoenolpyruvate from oxaloacetate in the BS cells by means of PEPCK (**Figure 2A**). The list of gene families that we considered as core or variable is found in (**Supplemental Table 4**).

**Figure 2:**
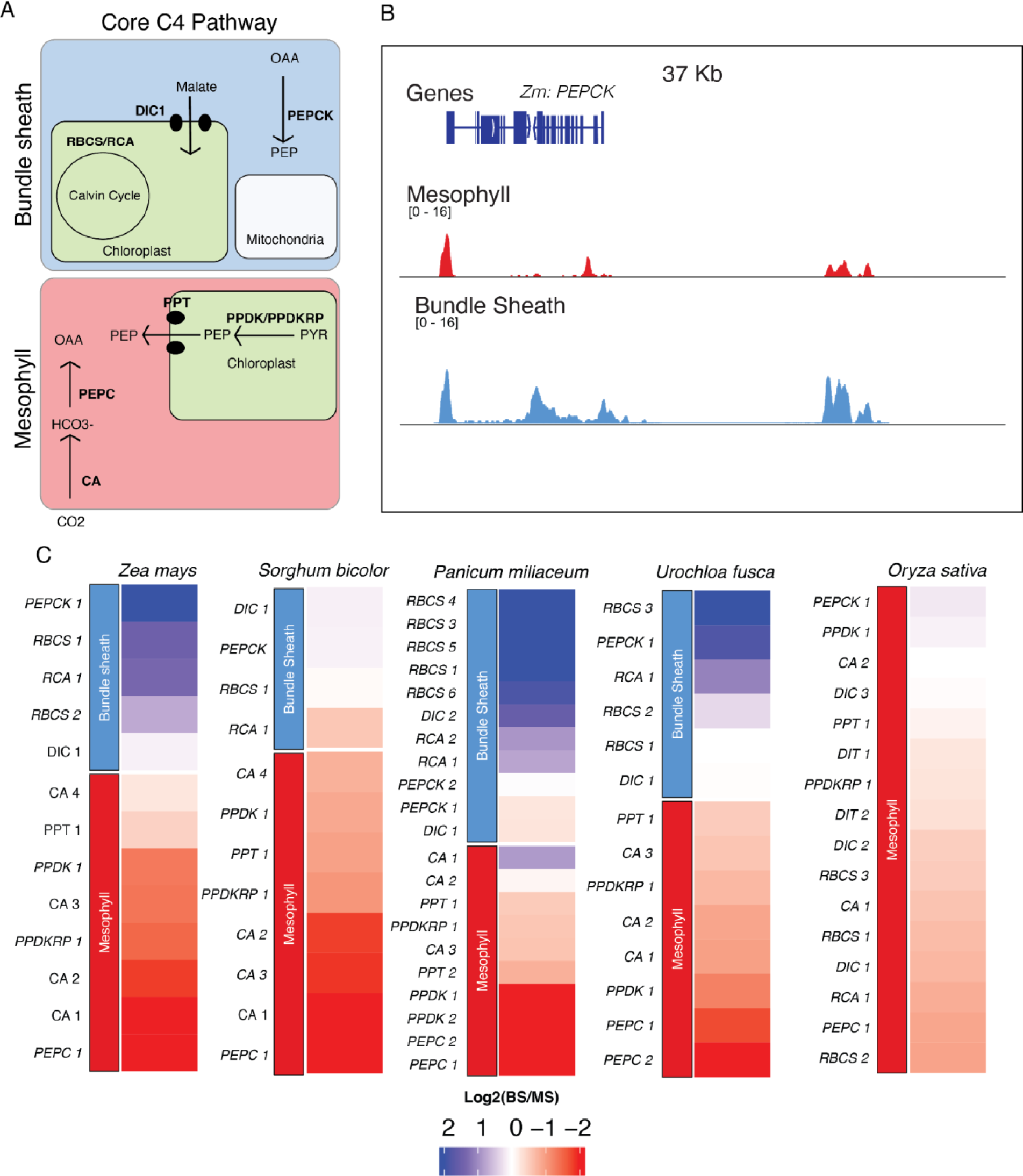
Cell-type chromatin-accessibility bias for core enzymes in C_4_ and C_3_ species. **A**) Schematic of the core C_4_ enzymatic pathway. Core C_4_ enzymes are defined as those which maintain their cell-type-specificity in all C_4_ subtypes sampled. The red and blue squares represent MS and BS cells, respectively. Enzymes are labeled in bold, and transporters are denoted by shapes. Intermediate molecules are indicated by non-bolded text. **B)** Screenshot of *PEPCK* in *Z. mays*. Blue tracks correspond to BS chromatin accessibility and red tracks show MS chromatin accessibility. Tracks are equally scaled to facilitate comparison. **C)** Heatmaps of chromatin accessibility bias of the core C_4_ enzymes. Values within each heatmap correspond to Log2(BS/MS). Blue indicates increased BS chromatin accessibility and red indicates increased MS chromatin accessibility. Each species column and subtype was clustered independently, and genes were assigned as being MS- or BS-specific (top/bottom of heatmap) based on literature. Enzyme copies were distinguished phylogenetically.

To investigate the cell-type bias of these enzymes, we used chromatin accessibility of the gene (gene body as well as 500 bp upstream of the transcriptional start site) (**Figure 2B**). Cell-type bias was calculated as the log_2_ fold change of BS/MS chromatin accessibility. To identify core C_4_ enzymes across these species, we used OrthoFinder, named and numbered the enzyme models based off of their relatedness to *Z. mays* copies of known core C_4_ genes (31). Using only cell-type-specific chromatin accessibility data, we observed expected cell-type bias with many orthologs of the maize MS-specific core C_4_ genes showing MS-specific bias as compared to BS (**Figure 2C**). For instance, in all C_4_ species, *PEPCK*, which regenerates PEP from OAA in BS cells, always showed a BS-specific bias (**Figure 2 A & C**). Additionally, *PEPC*, which converts bicarbonate to OAA in MS cells, showed MS-specific bias for all species sampled, except the C_3_ outgroup *O. sativa* (**Figure 2A & C**). These results highlight the quality of the data and the cell-type annotations for these single-cell datasets.

When analyzing these data in tandem with the phylogenetic trees, we noticed that some of the key enzymes showed different cell-type specificity based on their evolutionary origin (**Supplemental Figure 21-22**). For instance, for carbonic anhydrase in *P. miliaceum,* the orthologs that showed the largest bias between MS and BS cell types were not the copies that were the most evolutionary closely related to the *Z. mays* and *S. bicolor* cell-type-specific copies (Here *PmCA1* and *PmCA2*). Rather, a copy found in a separate clade (*PmCA3*) showed the most MS-specific bias (**Figure 2C**). This indicates that during the evolution of C_4_, different sets of carbonic anhydrases were likely co-opted for C_4_. One challenge using chromatin accessibility in this context, however, is the fact that neighboring gene models can occlude cell-type-specific signals. For instance, in the *S. bicolor* copy of *RBCS1*, a BS-specific gene has a neighboring gene model directly upstream which shares a promoter region making measurement of the cell-type-specific bias of some loci challenging when using chromatin accessibility data (**Supplemental Figure 23**).

One unexpected result from this analysis was the lack of cell-type-specific bias for *MALATE PHOSPHATE ANTIPORT 1 (DIC1),* also known as *DICARBOXYLATE/TRICARBOXYLATE TRANSPORTER 1* (*DTC1*) in *Z. mays*. It has been previously reported that *DIC1* had BS-specific expression bias in *Z. mays* as well as in *P. miliaceum* (32–34). However, there is not a clear signal based on the chromatin accessibility data. This could indicate that some ACRs harbor multiple CREs active in different cell types that are not obvious in chromatin accessibility data or that the cell-type-specificity observed is not due to *cis*-regulation, possibly involving post-transcriptional processes (**Figure 2C**). Lastly, as expected, there was very little bias in the C_3_ outgroup (*O. sativa*). In total, 12/13 of the core C_4_ enzymes showed cell-type-specific bias in *Z. mays*, 7/12 in *S. bicolor*, 16/21 in *P. miliaceum*, 11/13 in *U. fusca*, and finally 0/16 in *O. sativa*. These data demonstrate that chromatin-accessibility data can be leveraged to investigate the cell-type regulation of C_4_ genes while also taking into consideration their evolutionary relationships in a cross species context.

### Key C_4_ Subtype Enzymes Show Potential Convergent Evolution in Cell-type-specific Bias

We investigated the variable enzymes that give each C_4_ subtype its unique properties by focusing on two species (*S. bicolor* and *Z. mays*) from the *NADP-ME* subtype (**Figure 3A**). As expected, chromatin accessibility bias was observed for enzymes previously reported as having cell-type-specific expression patterns, similarly to the core C_4_ enzyme set (29,35). Reassuringly, one of the most biased enzymes identified was *NADP-ME*, the key enzyme of the redox step in *NADP-ME* subtypes. More specifically, of the multiple copies of *NADP-ME* that exist in *Z. mays,* we observed the expected cell-type bias for the known BS-specific copy, *ME3,* a key factor in C_4_ (here *ZmNADP-ME1*) (**Figure 3B**). We noticed in *S. bicolor*, the BS-specific *NADP-ME* and the MS-specific *NADP-malate dehydrogenase* (*NADP-MDH*) gene copies are recent tandem duplications, each maintaining their respective cell-type specific chromatin accessibility (**Figure 3B & C, Supplemental Figure 22**). The malate transporters *DICARBOXYLIC ACID TRANSPORTER1/2 (DIT1/2)* also demonstrated their expected cell-type-specific bias with *DIT1* being MS specific and *DIT2* being BS specific in both species (**Figure 3B & C**). However, upon further inspection of the phylogenies of the *DITs* in *S. bicolor,* we noticed a pattern where the most BS-biased copy, *SbDIT4 (*Sobic.004G035500), was phylogenetically more closely related to the *ZmDIT1*. Something which has been previously reported (33,36). These results indicate that over evolutionary time, even members of the same C_4_ photosynthetic subtype, which likely share a C_4_ ancestor, can use different paralogous loci to achieve cell-type-specific expression. This highlights that C_4_ evolution is an ongoing process.

**Figure 3:**
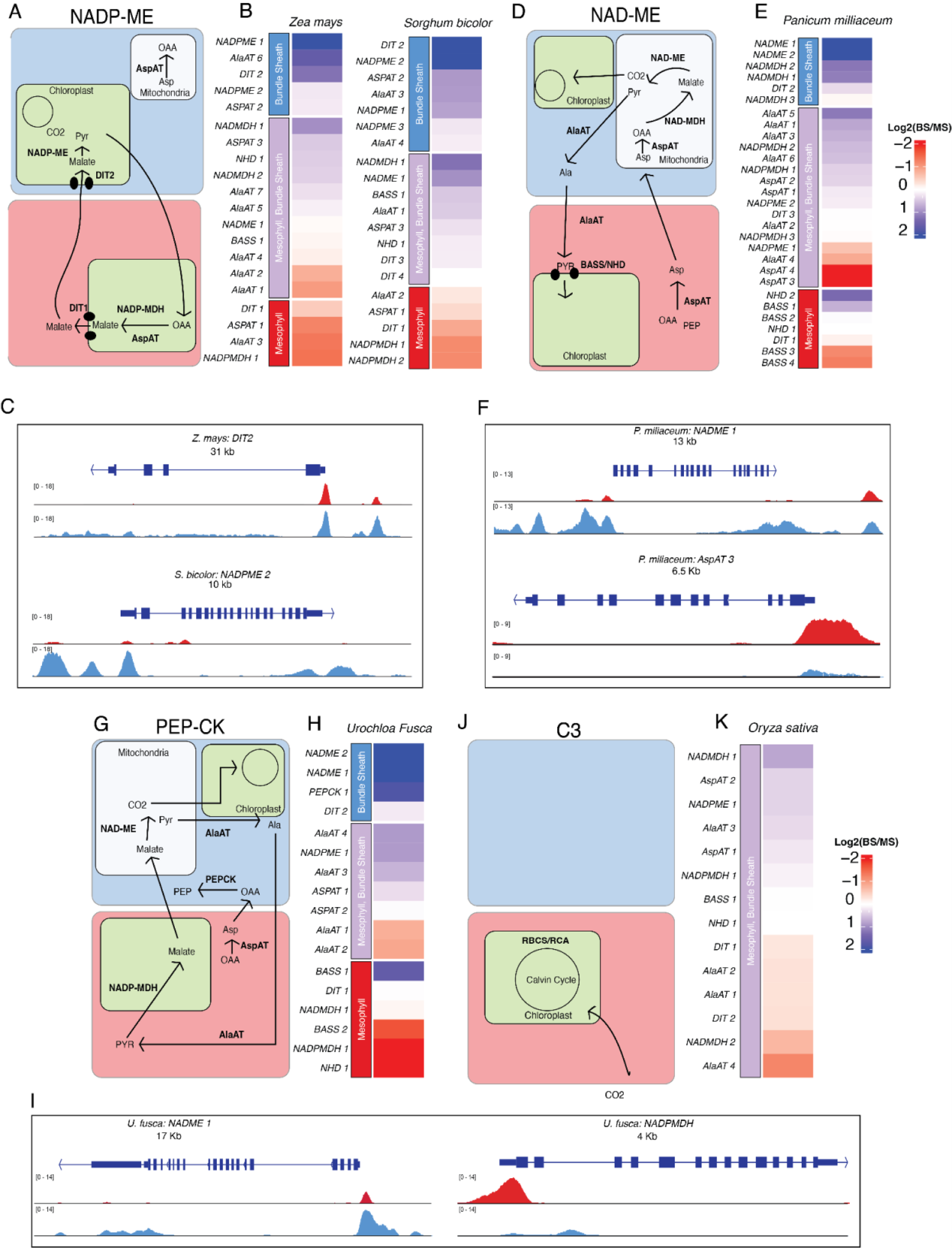
Cell-type chromatin accessibility bias for variable C_4_ genes associated with C_4_ subtypes. **A/D/G/J)** Schematic of C_4_ enzymatic pathways for various C_4_ subtypes. The red and blue squares represent MS and BS cells. Enzymes are labeled in bold, and transporters are denoted by shapes. Intermediate molecules are indicated by non-bolded text. For clarity, core enzymes have been removed. **B/E/H/K)** Heatmaps of chromatin accessibility bias in C_4_ subtype enzymes. Values within the heatmap correspond to Log2(BS/MS). Blue indicates increased BS-chromatin accessibility and red indicates increased MS-chromatin accessibility. Genes were labeled as being BS specific (blue) BS/MS specific (purple) or MS specific (red) based on previous literature. **C/F/I)** Screenshot of various C_4_ sub-type enzymes and their chromatin accessibility profiles around the TSS. Blue tracks correspond to BS chromatin accessibility and red tracks show MS chromatin accessibility. Tracks are equally scaled to facilitate comparison.

NAD-ME subtypes in *P. miliaceum* are interesting, as the intermediate molecule being passed between MS and BS doesn’t take the form of malate, but instead aspartate, alanine, and oxaloacetate (**Figure 3D**). At least one copy of all of the key redox enzymes, NAD-ME and the NAD-dependent malate dehydrogenase (NAD-MDH), show BS-biased chromatin accessibility (**Figure 3E & F**). Interestingly, of the three copies of *NAD-MDH* analyzed, only two showed bias for BS. Next, we evaluated two key enzymes associated with the generation of critical intermediate metabolites, Aspartate aminotransferase (AspAT), and Alanine aminotransferase (AlaAT). It has been reported that some AspAT have cell-type-specific expression patterns, with the MS-specific copy of the protein being transported to the cytosol and the BS-specific copy being transported to the mitochondria (**Figure 3E & F**) (37–39). Of the four copies of AspAT we examined, two (*PmAspAT3/4*) showed significant MS-specific bias, whereas the other two copies (*PmAspAT1/2*) didn’t show significant deviation towards BS (**Figure 3E**). This possibly indicates differing levels of regulation for the AspAT copies that did not show the expected BS bias, or missing copies of AspAT that we have not investigated. Within AlaAT, however, we identified one copy, *PmAlaAT1,* showing MS-specific bias, and *PmAlaAT6* showing BS-specific bias; something that has been previously hypothesized based on biochemical information (40). Additionally, somewhat unexpectedly is that we didn’t observe clear bias for sodium bile acid symporters (*BASS*) and sodium:hydrogen antiporters (*NHD*) (**Figure 3E**). These two proteins together form a functioning sodium bile acid symporter system, which balances the ratio of sodium and is important in the transport of pyruvate into the chloroplast of MS cells (41). Although two copies of the *BASS* genes were MS biased, only a single copy of *NHD* was slightly MS biased. Surprisingly, we do observe slight cell-type-specific chromatin accessibility bias for malate transporter *DIT1/DIT2* in *P. miliaceum*. This is somewhat surprising, as malate is not the main 4-carbon intermediate used by NAD-ME subtypes (10). This highlights the flexible nature of *P. miliaceum* in terms of its C_4_ photosynthetic style, as it has been implicated that it can perform some of the metabolite shuttling as the NADP-ME subtype (10,42,43). The potential flexibility of *P. miliaceum* in its style of C_4_ makes it an extremely interesting species to study, especially when considering that it doesn’t share common C_4_ ancestry with *Z. mays* or *S. bicolor.* This lack of evolutionary relationship between *P. miliaceum* and *S. bicolor* and *Z. mays* makes the comparison between *P.miliaceum* and its closer relative*U. fusca* all the more valuable. These observations point to the complicated nature of some of these C_4_ photosynthetic subtypes. While the obvious subtype-specific enzymes show expected chromatin-accessibility bias, others do not.

Using the *PEPCK* subtype in *U. fusca,* we evaluated cell-type bias of enzymes that operate as an intermediate between NAD-ME and NADP-ME subtypes (**Figure 3G**). Copies of *NAD-ME* and *PEPCK* showed significant BS bias (**Figure 3H & I**). Additionally, *NADP-MDH* was significantly biased towards MS, reflecting its critical role in the regeneration of malate from pyruvate (**Figure 3H**). We also observed one copy of *BASS,* which was heavily MS biased, as well as the only copy of *NHD* being highly MS biased (**Figure 3G**) (44). Within the *BASS* family, based on the phylogenies, it appears one clade of *BASS* genes was co-opted to be MS specific, whereas the other clade remained somewhat BS specific. This potentially indicates that this co-opted clade may have been predisposed for C_4_ photosynthesis at the common ancestor of *P. miliaceum* and *U. fusca.* Additionally, we also find one MS-biased and one BS-biased version of AlaAT (**Figure 3H**).

Finally, when evaluating genes in the C_3_ outgroup *O. sativa,* we only observed significant chromatin accessibility bias for three of the 14 enzymes. This is expected given the overall lack of enzymatic bias seen in C_3_ species (**Figure 3K**). Interestingly though, we did find a single instance where one copy of *AspAT* is BS specific, suggesting that this copy of *AspAT* might slowly be co-opted into being more BS-specific (**Figure 3K**). Even more interesting is the slight BS-specific bias of the rice *NAD-MDH*, a BS-specific enzyme in the *NAD-ME* subtypes. These results show a series of complex evolutionary relationships where many different genes can be co-opted into the C_4_ pathway, and highlights the myriad ways in which C_4_ evolution occurs.

### Cell-type-specific Accessible Chromatin Regions of Both Core- and Subtype-Specific Enzymes

Although measuring the gene body chromatin accessibility of C_4_ enzymes is valuable, it doesn’t inform us about the cell-type-specific *cis*-regulatory environment controlling these genes, as we only included 500 bp upstream in this initial analysis. To identify all potential CREs important for regulation of C_4_ enzymes, we identified cell-type-specific ACRs using a modified entropy metric (**Methods; Supplemental Figure 33-34**). In short, cell-type-specific ACRs are those which are unique to either a single cell-type or two or three cell-types in contrast to broadly accessible ACRs which are accessible in many different cell-types. For each C_4_ enzyme, in both the core and the non-core set, we identified ACRs around them. We only considered ACRs to be potential regulators of a locus based on distance, with assigned ACRs needing to be less than 200 kb away from the target enzyme, and requiring that no other gene intervenes between the ACR and enzyme in question. In total, across all variable and core enzymes and taking into consideration only C_4_ species, we find that on average, C_4_ genes have between 2-3 cell-type-specific ACRs, with an additional 2-3 broadly-accessible ACRs (**Figure 4A, Supplemental Table 5**).

**Figure 4:**
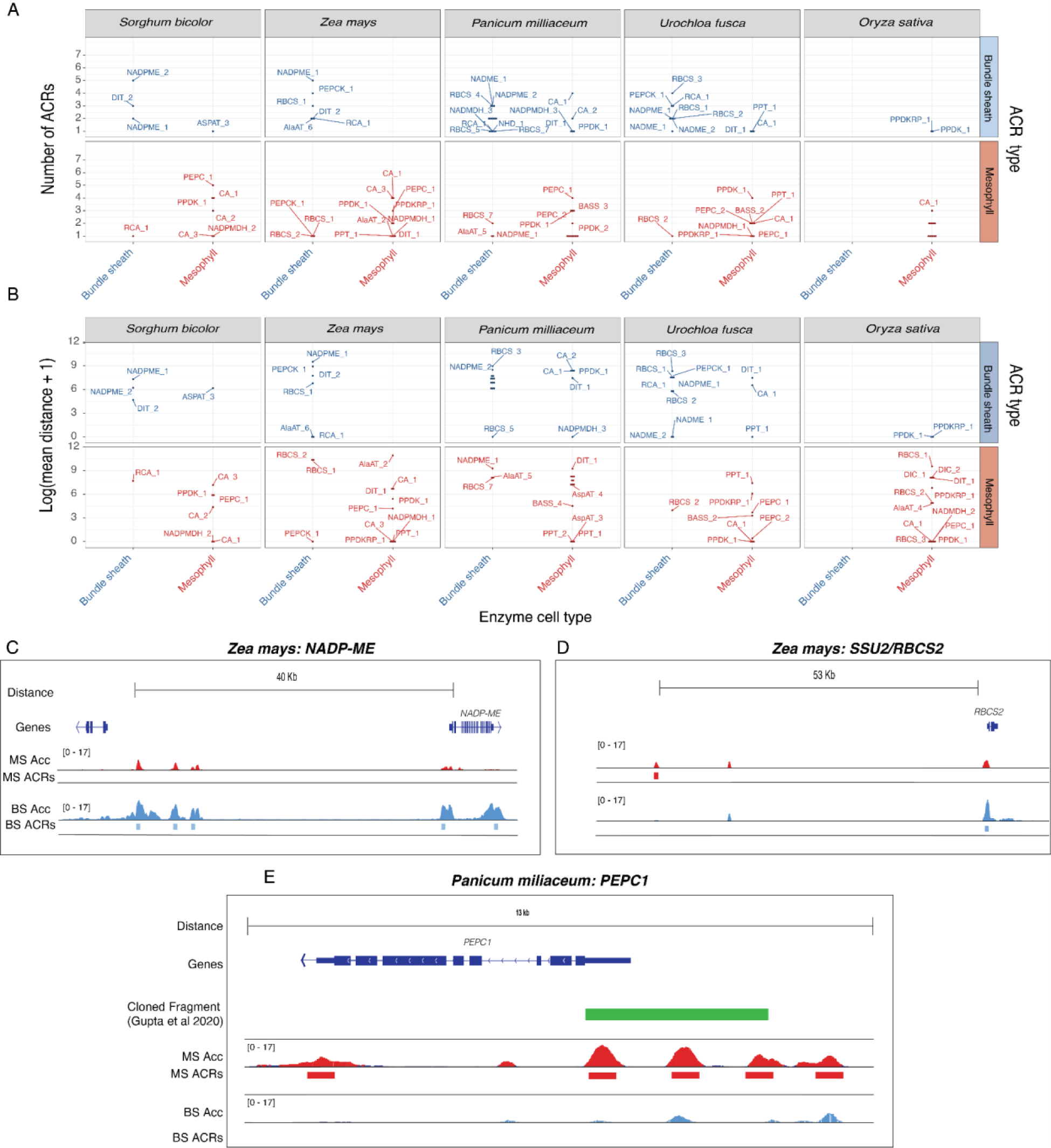
Investigating the number and distance of cell-type-specific ACRs around C_4_ enzymes across subtypes. **A)** Dot plots showing the number of cell-type-specific ACRs around each enzyme. The x-axis indicates which cell type these enzymes are found in. The y-axis is counts of ACRs. The graph is further subdivided with the top panel being broad ACRs, middle panel BS-specific ACRs, and the bottom being MS-specific ACRs. Enzymes are labeled. **B)** Dotplots showing the mean distance of cell-type-specific ACRs to their closest C_4_ enzyme. The x-axis indicates which cell type these enzymes are found in. The x-axis is the genomic distance to the C_4_ enzyme in question. If an enzyme had multiple cell-type-specific ACRs, the distance was averaged (mean). **C)** Screenshot of *NADP-ME1* in *Z. mays*. Blue tracks correspond to BS chromatin accessibility and red tracks show MS chromatin accessibility. Tracks are equally scaled to facilitate comparison. All genes found within this window are shown. **D)** Screenshot of *RBCS2* in *Z. mays*. Blue tracks correspond to BS chromatin accessibility and red tracks show MS chromatin accessibility. Tracks are equally scaled to facilitate comparison. All genes found within this window are shown. **E)** Screenshot of *PEPC1* in *P. miliaceum.* The green fragment represents the cloned promoter from Gupta et al 2020, which was identified by minimap2 alignment. Blue tracks correspond to BS chromatin accessibility and red tracks show MS chromatin accessibility. Tracks are equally scaled to facilitate comparisons.

For all C_4_ subtypes, the key redox enzymes all showed BS cell-type-specific ACRs, potentially identifying critical CREs for proper cell-type-specific expression. For instance, in *Z. mays, NADP-ME1* had five BS-specific ACRs, in *S. bicolor, NADP-ME2* had five BS-specific ACRs, in *P. miliaceum, NAD-ME1* had four BS-specific ACRs, and in *U. fusca, PEPCK,* had three BS-specific ACRs (**Figure 4 A & C**). Additionally, of the MS-specific enzymes, we consistently observed numerous cell-type-specific ACRs around the carbonic anhydrase family. On average, there were 3.5 MS-specific ACRs for each copy of carbonic anhydrase across all of the species. This likely reflects the fact that carbonic anhydrase is critical in the initial steps of C_4_, and also important in CO_2_ sensing (45). We also noticed an intriguing pattern where enzymes which were accessible in one cell type had cell-type-specific ACRs of the other cell type. For instance, around *RBCS2,* a BS-specific enzyme, we found a series of MS-specific ACRs (**Figure 4D**). On average, we found 2.5 BS-specific ACRs around *RBCS* and 1.5 MS-specific ACRs. This contrasting pattern was observed in key photosynthetic enzymes in all of the C_4_ subtypes. This likely indicates that some of these ACRs contain CREs that negatively regulate *RBCS* in MS, as cell-type-specific CRE usage has been implicated as being an important driver in proper compartmentalization (46,47). The identification of ACRs around key C_4_ enzymes provides a detailed map about potential *cis-*regulators of these loci, which provides the basis for future investigation into the direct function of each of these ACRs and how they might be altering transcription in multiple different ways. These results show that there are likely multiple ACRs important to cell-type specificity of these enzymes.

Traditionally, the field has focused on *cis-*regulation within a set distance from the transcriptional start site, often 1-2 kb, which is thought to generally encompass the promoter (48). However, we observed abundant distal cell-type-specific ACRs for many of these key genes (**Figure 4B**). For instance, the average distance of an ACR to its C_4_ enzyme is 10,080 bp (*Z. mays*), 3,017 bp (*S.bicolor*), 4,260 bp (*P. miliaceum*), 2,358 bp (*U. fusca*), and 4,730 bp (*O. sativa*), indicating that the *cis*-regulatory space for these enzymes is far greater than previously appreciated, where a majority of the focus in the literature is on putative promoters. To test this, we compared the identified ACRs to a series of previously reported cloned promoters. We found that for *Zea mays* and *Sorghum bicolor* the ACR space identified includes significantly more regions that are distal to their target gene (**Supplemental Figure 23C, Supplemental Table 6)** (25,49,50).

The genome of *Z. mays* emphasizes this point, as the subtype-specific enzyme *NADP-ME* has three cell-type-specific BS ACRs distal to the transcriptional start site, with the furthest being 34,336 bp away (**Figure 4C**). These distal ACRs provide critical regulatory loci to further investigate. Interestingly, we found some enzyme/ACR pairs with opposite cell-type-specificity (*i.e.* BS-specific enzyme, MS-specific ACR). Many of these ACRs were distally located. For example, in *Z. mays,* the MS-specific ACR of *RBCS* was 36,171 bp upstream (**Figure 4D**). When investigating ACRs around promoters, we were struck at how often cell-type-specific ACRs occurred outside of the bounds of previously analyzed promoters. For example, in *PEPC* in *P. miliaceum*, a recent analysis demonstrated that a series of conserved non-coding sequences found between species were able to drive MS expression (27). When we looked at chromatin accessibility data of the promoter fragment which was cloned from *PEPC*, we identified many MS-specific ACRs within the cloned fragment, but an additional one upstream. This results shows the advantage of using scATAC-seq data to identify candidate CREs for certain genes, removing the guesswork of cloning fragments to investigate and providing a detailed cell-type-specific regulatory map of the locus (**Figure 4E**). Thus, scATAC-seq greatly improves the search space of the active CREs potentially driving cell-type-specific gene expression patterns.

### The Evolutionary Relationships of ACRs Associated with C_4_ Genes is Complex and Variable

Next, we explored the evolutionary histories of these ACRs. Due to the fact that the C_4_ subtypes come from different radiation events, (with *Z. mays* and *S. bicolor* likely sharing a C_4_ ancestor and *U. fusca* and *P. miliaceum* sharing a different C_4_ ancestor), we were curious to evaluate if a majority of the ACR space around these genes were either novel, or shared among these species. We implemented a pairwise sequence based approach by identifying sequence conservation of ACRs between the study species using BLAST (**Methods)**. The majority of important C_4_ genes have both novel, and conserved ACRs. For example, PPDK, a MS-specific enzyme, shares ∼25% of its ACRs across all species examined including the *O. sativa* C_3_ outgroup (**Figure 5A**). Interestingly, *RUBISCO ACTIVASE (RCA)*, a critical enzyme in photosynthesis which removes inhibitory molecules from the RuBisCO active site, had novel ACRs in all of the C_4_ species examined, whereas *RCA* in the C_3_ species *O. sativa* shared one ACR with all of the C_4_ species. This might indicate that each of the C_4_ species gained regulatory sequences at *RCA* or that *O. sativa* might have lost them (**Figure 5A**). Focusing on NADP-ME revealed notable divergence in its associated ACRs, even among closely related species. For example, in *Z. mays*, two out of nine ACRs linked to *NADP-ME1* were unique, lacking counterparts in other species (**Figure 5A**). This is particularly striking given that *S. bicolor*, belonging to the same C_4_ subtype, diverged from *Z. mays* only 13 million years ago (51). Similarly, in *S. bicolor*, the BS-specific *NADP-ME2* variant exhibited two out of five unique ACRs. This pattern underscores the rapid and distinct evolutionary trajectories of ACRs in C_4_ plants. A full list of gene families, and gene models, and their relative conservation is found in **Supplemental Figure 25A.** Using this same approach to study all of the core class of C_4_ enzymes did not reveal a generalizable pattern associated with gain or loss of ACRs around C_4_ genes (**Supplemental Figure 25A**). Our findings not only confirm the dynamic evolution of *cis*-regulatory sequences in C_4_ enzymes but also align with existing research that highlights rapid *cis*-regulatory changes among closely related species (48,52).

**Figure 5:**
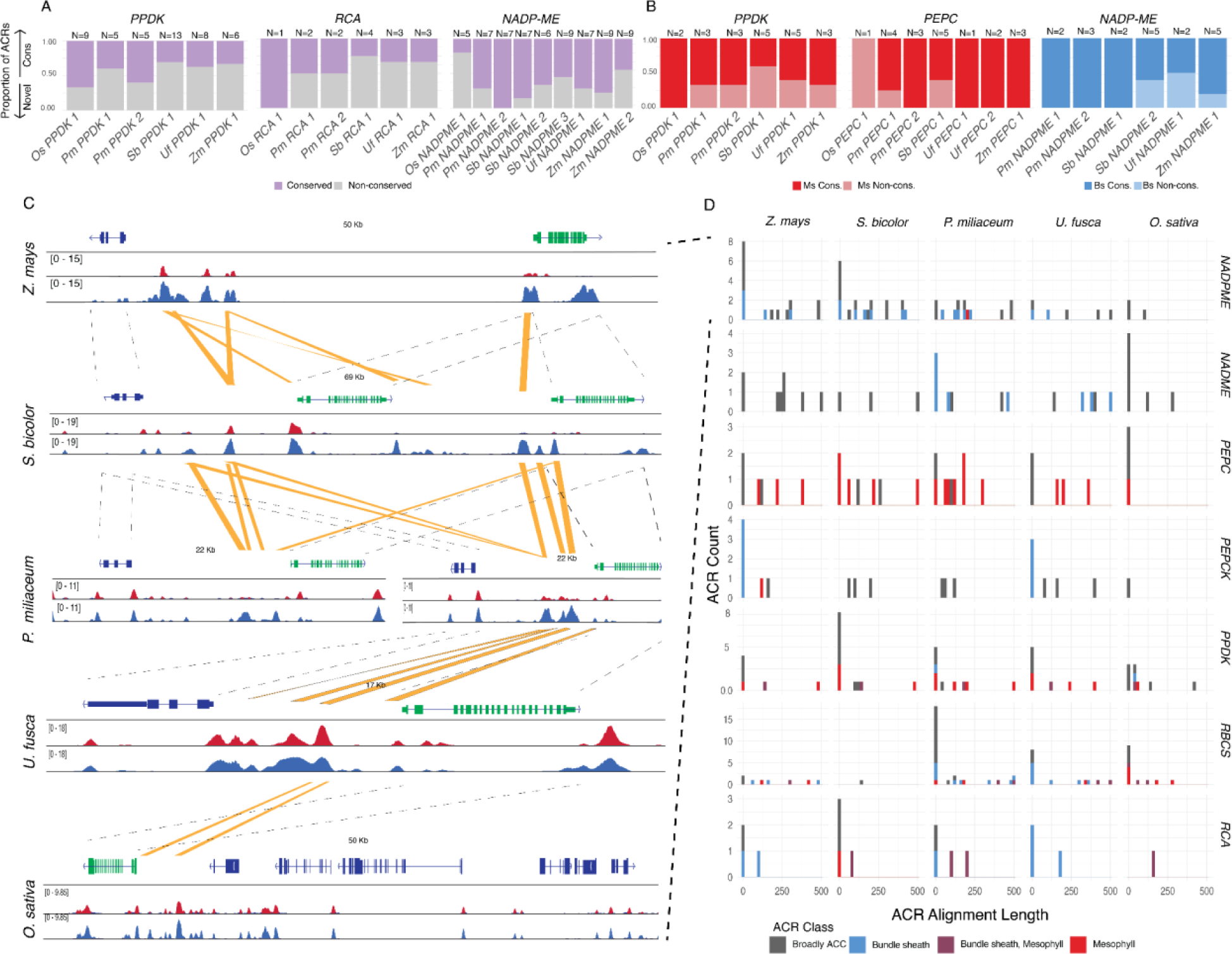
The evolutionary relationships of *cis-*regulatory regions around C4 genes is complex, being composed of both novel and conserved ACRs. **A)** The proportion of all ACRs that are conserved or novel for the following gene families *PPDK*, *RCA*, and *NADP-ME*. Purple bars represent ACRs that have any sequence aligned to them from a different species, and gray represents ACRs where sequences are not alignable. The number of ACRs in each locus is labeled at the top of each column. **B)** The proportion of cell-type-specific ACRs that are conserved and novel for the following gene families, *PPDK*, *PEPC*, and *NADP-ME*. Red bars only consider MS-specific ACRs, and blue bars only consider BS-specific ACRs. **C)** Screenshot of the conservation of BS-specific ACRs around *NADP-ME* across species. From top to bottom the species are *Z. mays, S. bicolor, P. miliaceum, U. fusca, and O. sativa. NADP-ME* is annotated in green for all species. Dashed bars between gene models represent the same gene model, and yellow bars are conserved ACRs. Browser tracks are blue for BS, and red for MS. Browser tracks are scaled within each species to allow for direct comparisons. **D)** The length of ACRs that are conserved in a cross species context. Rows represent gene families, and columns represent species. Each histogram is the number of ACRs within the loci of that gene family. The x-axis is the length of the ACR that is conserved and the y-axis is the count. ACRs are color coded according to the legend.

While investigating the ACRs around the C_4_ genes is interesting, understanding how cell-type specificity is achieved across C_4_ subtypes is needed for efforts to engineer C_4_ photosynthesis. When looking at just the cell-type-specific ACRs around key C_4_ loci, we find a similar pattern where there is a mix of both conserved and novel ACRs. For example, we discovered that some of the MS-specific ACRs associated with *PPDK* and *PEPC* are highly conserved in all of the studied species. Interestingly, the MS-specific ACRs around *PEPC* were only found in the C_4_ species, and not in the C_3_ outgroup, *O.sativa* (**Figure 5B**). This indicates that some of the CREs that allow *PEPC* expression in MS likely evolved after the split between the most recent common ancestors. We also observed that *NADP-ME* possessed numerous BS-specific ACRs that were conserved in all species, including *O. sativa* (**Figure 5B**). Considering the fact that proper compartmentalization of *NADP-ME* in BS cells is only critical in two of the three C_4_ subtypes, this was surprising. However, in both *S. bicolor* and *Z. mays*, there were novel BS-specific ACRs associated with each key *NADP-ME*. In *Z. mays,* one out of the five BS-specific ACRs was novel to *Z. mays*, and in *S. bicolor* two out of the five were novel to *S. bicolor.* Upon inspection of all the *NADP-ME* loci in genome browsers, we were struck by the complexities and shuffling that occurred at these BS cell-type-specific ACRs (**Figure 5C**). These results highlight that extensive *cis-*regulatory evolution is occurring in each of these species, and in particular on a cell-type-specific level. Additionally, this may point to the fact that the novel BS-specific ACRs found in *S. bicolor* and *Z. mays* may be more important for proper BS-specific expression than the conserved regulatory elements.

Although binary classification of ACRs was useful to decipher larger scale patterns between key enzymes, we next tested if larger segments of sequence were conserved around some C_4_ genes as compared to others. We profiled the relative amount of conserved sequence at each of these ACRs, as alignment of sequence between species gives greater resolution about important ACRs. One interesting observation from this analysis was the fact that the cell-type-specific ACRs around *PEPCK* appear to be novel between *Z. mays* and *U. fusca* (**Figure 5D, Supplemental Figure 29-30**). This suggests that these regulatory loci emerged independently, and yet are still likely important in cell-type-specific expression of *PEPCK*. Additionally, around the *NAD-ME* loci in *P. miliaceum*, we found diverse evolutionary histories with both copies *NAD-ME1* and *NAD-ME2* having both conserved and novel BS-specific ACRs (one out of four ACRs were novel for *NAD-ME1*, and zero out of the two were conserved for *NAD-ME2*) (**Figure 5D**). The ACRs from *NADP-ME1* are conserved in *U. fusca*, whereas all three BS-specific ACRs are conserved in relation to *P. miliaceum.* Pointing to the fact that the ACRs have likely maintained their cell-type specificity, and are likely critical drivers in the correct expression of *NAD-ME* loci. These results highlight the dynamic evolution of cell-type-specific ACRs around key C_4_ loci, and that even closely related subtypes have evolved novel ACRs potentially critical in terms of proper gene expression, as well as compartmentalization.

### Identification of *de novo* TF-Binding Motifs from Cell-type-specific Chromatin Data Reveals Rapid Sequence Diversification of ACRs

Leveraging the cell-type-resolved datasets, we identified *de novo* cell-type-specific TF motifs in BS and MS ACRs (**Figure 6 A & B; Methods; Supplemental Figure 31**). We selected the BS-specific motifs based on motif similarity within C_4_ species for BS, and motif similarity seen across all species for MS. Additionally for the identification of BS specific motifs, we identified motifs which didn’t appear to have a corresponding motif in *O. sativa* (**Methods**). Reassuringly, within the BS-specific motifs, we identified a DOF TF motif, which is a key driver in the switch to C_4_ photosynthesis (29,53,54). In brief, the DOF TFs have been implicated as being potential drivers of proper gene expression in *Z. mays* C_4_ genes, both as repressors and activators. For example, *ZmDOF30* has been implicated as being important in driving BS specific gene expression (29,53,54). In total we identified three BS-specific motifs, and four MS-specific *de novo* motifs that are shared between the species sampled (**Figure 6 A & B; Supplemental Figure 31**). Using motif comparison tools, we were able to assign five out the of the six motifs to a putative TF family, implicating potential novel regulators in BS-and MS-specific gene expression (**Methods; Supplemental Figure 29**). We surveyed the C_4_ ACRs for the presence and absence of these motifs to determine if they provide the information needed for cell-type specificity. We additionally overlaid our BLAST results from the previous analysis in order to explore the relationship between these motifs and conservation (**Figure 6C**). A substantial number of motifs were present within the non-conserved regions of the ACRs. For instance, in one MS-specific ACR associated with *ZmCA3*,12/13 MS-specific motifs were found in non-conserved regions, suggesting these regions could be critical for driving the cell-type-specificity of this locus (**Figure 6D**).

**Figure 6:**
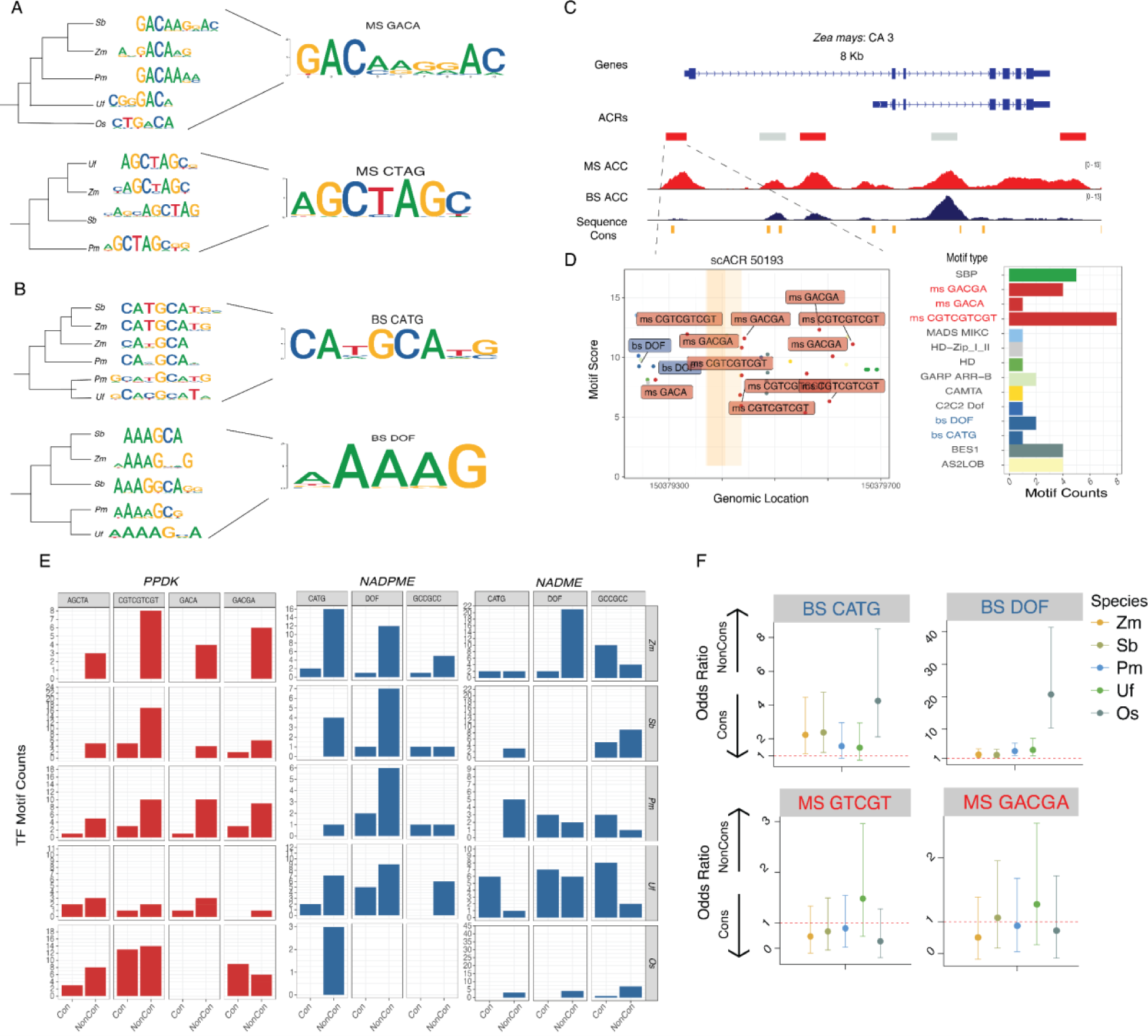
Identification of cell-type-specific TF motifs reveal a complex relationship between sequence conservation and motif prescnece. A subsample of MS-(**A**) and BS-specific (**B**) *de novo* TF motifs identified. **Left)** *De novo* motifs were clustered by the correlation of their PWMs and a correlation based tree was generated. **Right)** Representative PWMs from *de novo* discovery. **C)** Screenshot of the *ZmCA3* locus. ACRs are color coded based on their cell-type specificity. MS- and BS-chromatin accessibility tracks are equally scaled for comparison. Sequence conservation is identified by the ACR having sequence homology to other *CA* ACRs from a different species. **D)** An example of the conservation and motif landscape of one MS-specific ACR at *ZmCA3*. Left, the location of the motifs in ACRs with MS- and BS-specific motifs labeled. Orange highlighted regions correspond to the region of sequence conservation seen above. Right, quantification of the motifs found in the ACR. X-axis is the motif count, and the y-axis is the motif. **E)** The counts of TF motifs in conserved and non-conserved ACRs for three different genes across all five species. Y-axis is the number of ACRs of a given type, and the x-axis indicates the type of ACR. **F)** Odds ratio of four motifs when comparing their enrichment in conserved versus non-conserved regions. A higher odds ratio indicates that the motif is more often found in non-conserved regions within ACRs, whereas a lower odds ratio means the motif is in conserved regions. The cell-type-specific motifs found in **A/B** are colored in red and blue, respectively.

We expanded the analysis of BS- and MS-specific motifs in conserved and non-conserved regions of ACRs across key loci in the C_4_ species. On average the MS-specific motifs are more conserved than the BS-specific motifs (**Figure 6E-F; Supplemental Figure 32**). Agreeing with previous models of C_4_ evolution where some motifs that are MS specific have been co-opted to operate in C_4_ photosynthesis (**Figure 6D**) (11). Interestingly, we noticed a pattern where around *PPDK*, many of the MS-specific motifs appeared to be in non-conserved sequences for all of our species sampled (**Figure 6E**). This pattern is further highlighted in both *NADPME*, and *NADME* loci, where a majority of the BS-specific motifs occurred in non-conserved ACR regions for *NADPME*. This pattern is more nuanced in the *NADME* ACRs, as *P. miliaceum* and *U. fusca* share a significant amount of conserved sequence containing BS-specific motifs in the ACRs, suggesting that the BS-specific regulatory changes associated with these motifs are important (**Figure 6F**). These results highlight the capacity of genome-wide single-cell *cis*-regulatory maps to pinpoint key TF motifs important for the evolution of cell-type specificity.

### The DITs in the NADP-ME Subtypes Demonstrate Dynamic CRE Evolution

Upon analyzing the malate transporters *DICARBOXYLIC ACID TRANSPORTER’s (DITs* also known as the *DCTs)* we noticed the DITs in the NADP-ME subtypes showed an interesting pattern where the copies of *DIT1* in *Z. mays* and *S. bicolor* showed MS-specific chromatin accessibility, but the BS-specific copies of the *DITs* showed a more complex evolutionary history (**Figure 3B; Figure 7A**). We generated a phylogeny with additional species, and found that the BS-specific copy of *ZmDIT2* is related to two additional copies of *DITs* which are not BS-specific in *S. bicolor* (Here *SbDIT2.2* and *SbDIT2.1*) (**Figure 7A**). *S. bicolor* has a BS-specific copy of *SbDIT4,* which shares a clade with *ZmDIT1*. These results are consistent with earlier studies that found similar patterns and gene expression profiles of these copies of the *DITs* in *Z. mays* and *S. bicolor* (33,36,55). Although previous studies have documented changes in cell-type-specific gene expression for the BS-specific copies of the DITs, the mechanisms underlying these changes remain unclear. By using cell-type-specific ACRs, we explored if expression changes are associated with changes in the number of cell-type-specific cis-regulatory elements over evolutionary time.

**Figure 7:**
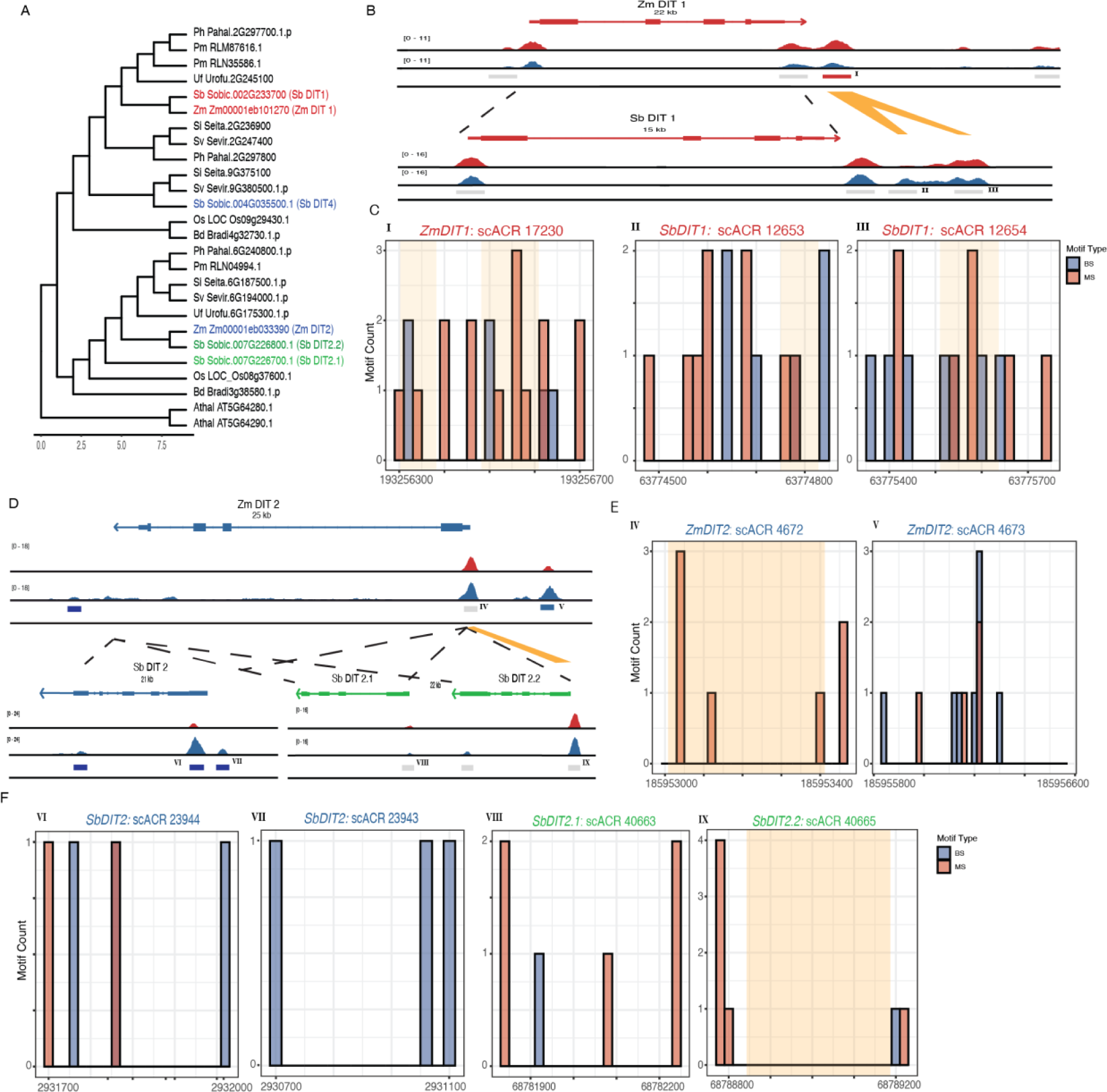
**A)** Phylogenetic tree showing the evolutionary relationship of the *DITs* in the monocots. *DITs* for *Z. mays* and *S. bicolor* are colored by their observed cell-type specificity, with red being MS specific, and blue being BS specific. Additional species have been added to increase resolution **B)** A screenshot of the *DIT1* between *Z. mays* (top) and *S. bicolor* (bottom). Yellow boxes indicate ACR sequences with conserved homology **C/E/F)** Motif location of BS and MS specific motifs in each ACR. The x-axis is the location within the ACR, and the y-axis is the motif count. Yellow bars indicate regions of sequence homology. Within each histogram, the x-axis is binned into 20bp regions for ease of graphing. Roman numerals in the top corner highlight the corresponding ACR found in the screenshot. **(I-IX) top)** X-axis the genomic coordinates of the given ACR. Yellow blocks denote the sequence homology as seen above. Y-axis, the motif score as calculated by motifmatchR, higher scores indicate a more confident motif. **bottom)** The count of each motif identified in the ACR. Note that BS and MS *de-novo* identified motifs are in blue and red respectively. **D)** A screenshot of the BS specific DITs loci between *Z. mays* (top) and *S. bicolor* (bottom). For the *S. bicolor* versions of the *DIT*s, *DIT4* is colored blue for its observed BS specificity and *DIT2.1* and *DIT2.2* are colored green. Yellow boxes indicate sequence homology.

To understand how cell-type specificity changed in these DITs due to changes in *cis*-regulation, we compared the ACRs associated with the *DITs*, and mapped the TF-binding motifs found within each ACR (**Methods**). For the MS-specific *DIT1s*, we focused on a MS-specific ACR located at the 3’ end of *DIT1* in *Z. mays* (**Figure 7B**). Upon comparing this ACR to *S. bicolor*, we were struck that the sequence found in the *Z. mays* ACR was actually split in two in *S. bicolor*, neither of which demonstrated cell-type specificity in *S. bicolor* (**Figure 7B; Supplemental Figure 33**). A closer inspection of motifs in these ACRs showed many MS-specific motifs (**Figure 7B-C**). These motifs might promote MS-specific gene expression of this locus. However, many *S. bicolor* MS-specific ACRs were not found in regions with any homology to *Z. mays* (**Figure 7C**). These results point to the rapid change of candidate CREs (cCRES) in this locus, and likely indicate that cCREs important in cell-type-specific gene expression might not be only found in conserved regulatory regions (56). Rather, selection of MS-specific gene expression is ongoing, and may yield significantly different regulatory environments in relatively short evolutionary time scales.

Next, we examined the BS-specific *ZmDIT2* and its two orthologs *SbDIT2.1 and SbDIT2.2*, which are not BS specific (**Figure 7A, D**). The BS-specific ACR around *ZmDIT2* has many DOF TF motifs (**Figure 7E**). These motifs are interesting, as expression changes within the DOF TF family could be important in driving BS-specific gene expression in C_4_ plants (29,53,57). When comparing the BS-specific ACRs around *ZmDIT4* to the more closely related copies of *SbDIT2.1* and *SbDIT2.2,* we found no conservation of these DOF TF motifs, and rather a significant lack of BS-specific TF motifs (**Figure 7F**). Considering the fact that neither of these *DIT* copies in *S. bicolor* show BS-specific expression, this result makes sense. Potentially providing a model where the *ZmDIT4* locus either gained these cCREs allowing for this copy of *ZmDIT2* to have BS specific gene expression, or *S. bicolor* lost these BS-specific motifs, and had a gain in *SbDIT4* specificity. In either scenario, it demonstrates the rapid pace of CRE evolution, and how these regions might be altering cell-type-specific gene expression. These results are in contrast to *SbDIT4*, where the ACRs around this locus are BS specific, and contain BS-specific motifs identified in our previous analysis (**Figure 7F**). In total, these results highlight the rapid rate of regulatory change around key C_4_ loci, and highlight the fact that there are likely key regulatory switches outside of conserved sequences. Finally, these results emphasize the fast pace in which cell-type specificity changes in plants

## Discussion

Understanding the evolution of *cis*-regulation associated with C_4_ photosynthesis has been a long standing goal in the field of plant biology. In this study, we demonstrated the utility of single-cell ATAC-seq data to investigate many aspects of the evolution of C_4_ photosynthesis. By identifying cell-type-specific chromatin accessibility from four C_4_ species composed of three different C_4_ subtypes, as well as a single C_3_ outgroup, we were able to compare and contrast key genes and their ACRs which define and distinguish C_4_ photosynthesis. We have shown that by using gene-body chromatin accessibility data, we can measure cell-type-specific bias of both core, and subtype-specific C_4_ enzymes. When taken into consideration with the gene family trees of many of these enzymes, we show diverse co-option of enzymes into the C_4_ pathway. Additionally, we identify cell-type-specific ACRs surrounding these key C_4_ enzymes. We find numerous cell-type-specific ACRs surrounding key C_4_ enzymes, many of which fall outside of the core promoter region. Additionally we find that around all of the C_4_ enzymes there is a mix of both conserved and novel cell-type-specific ACRs indicating that regulatory evolution of these regions is ongoing. Finally, we use cell-type-specific ACRs to identify a series of *de-novo* binding motifs which appear to be cell-type specific, and show that these motifs surround C_4_ loci, and have a mixed relationship with conservation depending on the motif. This indicates that cell-type-specific TF motifs are rapidly changing around C_4_ loci.

Investigation of the CREs driving cell-type-specific expression of C_4_ genes is challenging. This often requires evaluation using transgenic plants, which limits the number of CREs that can be tested. This has greatly hampered efforts at understanding how *cis*-regulation of C_4_ genes evolves, whether by co-option of existing CREs or emergence of new ones. Our results show the complex nature of CRE evolution of C_4_ genes, including those specific to C_4_ subtypes. While we observe conservation of ACRs around many C_4_ genes, we do see interesting examples where the subtype-specific enzymes have evolved novel ACRs (*NAD-ME*’s in *P. miliaceum*, and *PEPCK* in *U. fusca*). These results support that there is likely a combination of both co-opting pre-existing CREs, as well as evolving new ones to facilitate proper expression and cell-type-specification of genes. This is further exemplified by the analysis of the *DIT* family of transporters, where we show striking accumulation of cell-type-specific TF motifs in non-conserved regions of ACRs between two closely related species. This highlights that the regions of the genome promoting cell-type-specific gene expression are likely found in both conserved, and novel regions. Another recent single-cell genomic study of the evolution of CREs important for photosynthesis using a comparison between *O. sativa* and *S. bicolor* reached similar conclusions (57). They frequently found different ACRs and TF motifs in promoters of orthologous C_4_ genes (57). Future efforts to assay these candidate CREs using reporter assays, transgenesis and genome editing will be required. Additionally, expanding these analysis outward to all genes associated with photosynthesis might provide valuable insights into how genes in the Calvin-Benson cycle alter their regulation in their adaptation to C_4_ photosynthesis. Fortunately, these high-resolution maps of cell-type-specific ACRs of these key genes/species provide a strong foundation to build upon.

Although these studies provide a blueprint for the study of key candidate CREs associated with C_4_ enzymes, profiling cell-type-specific chromatin accessibility of additional species would be greatly beneficial. Although *O. sativa* is an invaluable outgroup for this study, additional more closely related C_3_ species might make these comparisons simpler, and add additional resolution. For instance the C_3_ grass species *Dichanthelium oligosanthes* is more closely related to *U. fusca* and *P. miliaceum* and has a recently completed reference genome (58). Adding more species would enable greater resolution in the comparison of cell-type-specific ACRs, as the genetic distance between the species we examined and *O. sativa* make identification of conserved and novel ACRs challenging. As an example, the ACRs associated with *NAD-ME’s* in *P. miliaceum* might be co-opted instead of novel, however, based on our sampling, we cannot say.

Genome editing analysis of many of these ACRs would significantly advance which ACRs, and more specifically which CREs within the ACRs are most important for cell-type-specific expression (22). However, currently generating genome edits in monocots is challenging, time consuming and expensive. Fortunately, improvements to transgenesis are constantly improving making achieving these goals more likely in the future (59). It’s also important to consider that mutational analysis of CREs is not straightforward, often requiring numerous editing events of the *cis*-regulatory landscape of each gene. Previous studies have shown that deletions of many CREs produce variable molecular and morphological phenotypes, further complicating our understanding of the *cis*-reglatory code (60–62). And finally, many species, including *P. miliaceum* and *U. fusca* have to date never been transformed. This highlights the need to continually improve transgenesis methods to help facilitate the molecular dissection of CRE. In conclusion, this study provides a comprehensive map of cell-type-specific ACRs around key C_4_ genes, which reveals the dynamic evolution and diversity of *cis*-regulation of C_4_ genes.

## Acknowledgments

This research was funded by awards from the National Science Foundation (IOS-2134912 and IOS-1856627) and the Office of Research to RJS and Hong Kong University Grant Council (GRF 1409420) to SZ. APM and JPM were supported by the National Institutes of Health (K99GM144742) and (T32GM142623), respectively. This research was additionally funded with support from the NSFC (32100438 & 32370247) and Shanghai Jiao Tong University 2030 Initiative (to X.T.).

## Methods

### Plant Growth Conditions and Sampling

Seedlings of all five plant species, including maize (*Zea mays* B73), sorghum (*Sorghum bicolor* BTx623), proso millet (*Panicum miliaceum* L. CGRIS 00000390), and browntop signalgrass (*Urochloa fusca* LBJWC-52), along with the C_3_ plant rice (*Oryza sativa* Nipponbare), were grown under the conditions of 12:12 Light/Dark cycles at 30°C Light/22°C Dark and at 50% humidity.

The sampling of the C_4_ species was timed to coincide with a specific developmental stage, identified when the ligule of the third leaf became visible, marking the third leaf unfolding, yet prior to the appearance of the fourth leaf. For the C_3_ species, rice, 18-day-old leaves were used to correspond with the equivalent stage of the C_4_ species.

### Library Preparation

Nuclei isolation for the experiments was conducted using fresh seedlings of both the C_4_ and C_3_ species at their respective developmental stages. The methodology for nuclei extraction, encompassing the buffer composition and the subsequent steps, was used with procedures outlined for single-nucleus combinatorial indexing with transposed-based ATAC-seq library construction, as detailed in a prior study (63).

### Genomes

The *Z. mays* genome version 5 was downloaded from MaizeGDB (64,65). The *O. sativa* genome was downloaded from rice.uga.edu. The *S. bicolor* version v5.1 was downloaded and used from Phytozome version 13, as well as the *U. fusca* genome version 1.1 (66). Finally the *P. miliaceum* genome was downloaded from NCBI, bioproject number PRJNA431363 (43).

### Barcode Correction Read Alignment and Mapping of Tn5 Insertions

Read UMIs were processed using cutadapt (version 4.5) to identify UMIs (67). First, the index adapter sequences were trimmed from the reads. Next, the well barcodes and Tn5 barcode within the reads were identified, removed from the original sequencing read, and appended to the read header. Finally, a shell script is used to integrate all barcode information from the reads’ headers and label them correspondingly in the paired-end sequencing fastq files. Reads were aligned using BWA (version 0.7.17) (68). Reads were filtered using samtools (version 1.16.1) for mapping quality of >10 for *Z. mays*, *S. bicolor*, *U. fusca*, and *O. sativa*. *P. miliaceum* required a greater threshold of 30 given its recent whole genome duplication event increasing the rate of multi-mapping reads (69). Duplicate reads were removed using picard tools (version 2.25.0) (70). Single-base pair Tn5 integration events were mapped using the python script ‘makeTn5bed.py’ found in the GitHub utils directory (https://github.com/Jome0169/Mendieta.C4_manuscript). Finally, for each barcode only unique Tn5 integrations sites were used for analysis. So if a nuclei had the same identical fragments multiple times, only a single event was considered.

### Isolating High-Quality Cells

Cells were filtered using Socrates (21). In short, Fraction of Reads in Peaks (FRiP) scores were calculated for each cell by pseudo bulking the libraries and identifying peaks. For each individual cell, FRiP was calculated by intersecting Tn5 integration events with peaks. Cells with a FRiP score greater than 0.2 were used. Additionally, TSS enrichment was calculated by looking at the number of Tn5 integrations around TSS. Cells that had a TSS enrichment greater than 0.15 were used. Finally, cells were compared to a random sample of low quality cells which did not pass filtering, representing the “background” of cells, and correlation was calculated between passing cells and background cells using the corr package in R. Cells which had a correlation lower than 0.3 percent as compared to background cells were used for further analysis.

UMAP embeddings were then calculated for each species utilizing genomic bins (71). For each dataset, bins of 500 bp were calculated. To reduce the size of features to cluster on, bins had to show accessible chromatin in at least 0.005% of total cells (roughly 50∼100 cells in each species). Additionally, bins that were broadly accessible across greater than 10% of cells in the given dataset were also discarded to remove regions of the genome which were constitutively accessible and wouldn’t facilitate clustering. Finally, regions of the genome which were associated with either blacklist (21), or genes which were known to be related to cell cycle and circadian rhythms were removed. The final resulting matrix, which represented cell barcodes X genomic regions (here bins), were then put through the term-frequency inverse-document-frequency (TF-IDF) algorithm to identify genomic regions more descriptive of the entire dataset (30). The resulting matrix was then input into Singular Value Decomposition, and clustering was then done on the remaining features with the number of principal components (PCs) equaling 50, and any PC with a correlation to read depth greater than 0.5 removed (72) (30). Clustering was done using the Louvain clustering algorithm in order to bin cells into similar groups based off of the PCs calculated above, with parameters “res = 1.5, k.near = 30, m.dist = .01” in order to set K nearest neighbors to 30, minimum louvain distance to .01 in euclidean space (73). Using the UMAP embeddings, doublets were removed using the software Scrublet as implemented in Socrates software (74). At random, 5,000 cells were used to generate *in-silico* doublets, and cells which were scored as being likely doublets were removed. Adaptive thresholds were set on a per library basis. The doublet rate from Scrublet was compared against a mixed library where genotypes of *Z. mays* were mixed Mo17 and B73, and genotype doublets were identified. We found that Scrublet, on average, removed more cells in a conservative fashion than the birthday problem and genotype doublets identified, so we utilized the Scrublet doublet scores to be conservative. For the P*. miliaceum* dataset, replicates were found to integrate poorly in the UMAP embedding. Harmony (version 0.1.1) was used adjust replicate overlap with parameters “theta = 2, nclust=4, and var = “sampleID” (75). After integration, clusters which skewed greater than 75% towards one replicate were removed from downstream analysis.

### Identification of Putative Orthologs

To annotate species with less marker gene information, we identified putative orthologs or marker genes using OrthoFinder (version 2.5.4) (31). For each species, the primary protein sequence of the transcript was used as input to Orthofinder. In the resulting orthofinder outputs, the script “find_markers.orthofinder.py” was used to parse the resulting phylogenies and return back putative orthologs (https://github.com/Jome0169/Mendieta.C4_manuscript). For all C_4_ genes analyzed, each orthogroup was additionally annotated by hand in order to ensure accurate assignment of nearest orthologs phylogenetically.

### Annotation of Cell Types

Cell types were annotated by calculating gene chromatin accessibility for marker genes in each genome on a per cell basis. These values were then visualized on the UMAP embedding, and clusters with numerous marker genes associated with the same cell-type were used as evidence. Additionally, for each louvain cluster, enrichment of marker genes was calculated by comparing the cluster average as compared to a random shuffle of random cells. The top five most enriched markers were used in tandem with the UMAPs to ascertain cell-type identity. We also tested the statistical significance of the marker gene using Presto, a modified Wilcoxon rank-sum test in order to identify the most unique marker gene in each cluster (76). Additionally, for specific clusters showing mixed signals from marker genes, sub-clustering was done by isolating the cluster in question, and then re-clustering these cells on a new UMAP manifold. The same steps were done to visualize marker genes, as well as test this enrichment, and statistical significance. Finally, to bolster our set of marker genes across species, we used our most confident cell-type annotation in *Z. mays* to *de novo* discover marker genes. To do so, we utilized our gene-body-accessability metrics for each annotated cell-type, and ran DESeq2 (version 1.42.0) in a replicate aware fashion using all other cells as a null (77). Only statistically significant markers were kept which had a fold change greater than 1.5, and a log fold standard error of less than .6. OrthoFinder was used as mentioned above to find orthologs. To ensure that we were comparing similar cell-types, we also took an orthogonal approach where we compared the gene accessibility of the top 2000 most variable orthologs between our species. A linear model was used for each species comparison where the mean gene accessibility was taken into consideration, and the species was one-hot-encoded. Variation was calculated as the average variation between both datasets. The resulting residuals were used to generate the cell-type correlations.

### Peak Identification

To identify peaks, cells of the same annotation type were pseudo bulked in a replicate aware fashion. Within each replicate MACS2 (version 2.2.9.1) was run with parameters “--nomodel -- keep-dup auto --extsize 150 --shift −75 --qvalue .05” and variable genome size flag ‘-g’ (78). Summits for each peak identified in each replicate were extended by 250 bp in either direction. Only peaks which overlapped between replicates were used. To merge peaks from various cell types and select peak boundaries, the p-value associated with each peak in each cell type was compared by calculating the chromatin accessibility score for each peak per million, with those peaks with the highest accessibility score being selected as the representative peak. This method of identifying the most representative peaks across cell-types was inspired by previous single cell ATAC-seq papers (30,79,80). Additionally, bigwigs were generated for each cell type by normalizing each dataset to the number of reads/per million scaling factor. Implementation of this algorithm is found in the script call_scACRs.py for ease of use and replication in other experiments.

### Identifying Cell-type-specific ACRs

To identify cell-type specific ACRs, a modified bootstrapping method was used which drew inspiration from the modified entropy metrics found in (79). On a per ACR basis, Tn5 integrations per cell-type were summed and counts per million (CPM) normalized. These values were then converted to a probability by using the following equation (below, equation 1) where *pi* is the CPM value for the focal cell-type and *qi* is the total sum of all CPMs. From this probability statement, a modified shannon entropy metric was calculated, followed by a metric of specificity *Qpt*. For robust cell-type-specific ACR identification, the annotated cell-type was bootstrapped 5000 times, taking a sample of 250 cells from the cell population in question, and calculating both entropy and specificity scores. This was done to attempt to get a robust signal of specificity, which takes into consideration the variation in cell quality present in each cell-type annotation. To generate the null distribution of specificity scores, individual cell annotations were scrambled to generate an equal number of null cell-type classifications. For each null value, the entropy and specificity score were calculated. Finally to calculate a p-value, a non-parametric approach was used to identify how many of the real bootstraps fell outside of the null distribution using a one tailed test. ACRs which had a p-value of <0.001 were considered to be significant. ACRs were finally classified by the number of cell types they were specific to. ACRs specific to greater than three were classified as broadly accessible, less than or equal to three as cell type restricted, and a single cell-type as cell-type specific.

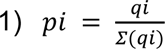

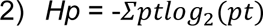

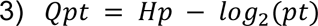

### Identifying Conserved ACRs Across Species

Since a majority of the C4 genes identified were not in synteny with one another, we took a gene family based approach to identify conserved and non-conserved ACRs associated with our C4 genes. In short, all ACRs within two gene models of a C4 gene are utilized for comparison. Sequences from the ACR were isolated using “bedtools getfasta” (version 2.31) (81). Then in a pairwise fashion each species had their ACRs from one C4 gene family compared to the corresponding genomic loci of the same gene family in a different species. Comparisons were made using Blastn (version 2.2.29) with the following parameters “ -task blastn-short -evalue 1e-3 -max_target_seqs 4-word_size 7 -gapopen 5 -gapextend 2 -penalty −1 -reward 1 -dust no -outfmt 6” (82). The output blast files were further filtered requiring sequence alignment to be greater than 20 nts, and have an evalue of .001. This analysis and the detailed commands ran can be found in the following snakemake file titled “ID_syntenic_orthologous.ACRs.snake”, and found in the snakemake directory in the associated github.

### Identifying Cell-type-specific Motifs

*De-novo* cell-type-specific motifs were identified by using XSTREME (version 5.5.3) of the MEME suite (version 5.5.5) package (83,84). In brief the sequences underlying the cell-type-specific ACRs were isolated, and equally matched null set of broadly-accessible ACRs were used the comparison for genomic enrichment. These null ACRs were matched in terms of GC content, and were only allowed to be 5% different from the cell-type-specific set in question and generated using the script “gen_null_fa.py”. Upon generation, motifs were analyzed using the universalmotifs package in R (version 3.18) (85). Motifs were first compared using HELL distance, and motifs which had a low correlation were discarded. In order to generate representative motifs, highly correlated motifs were merged using the function “merge_motifs” in found in the universalmotifs package. To identify the location of motifs, the R package motifmatchR were used, with a significant value cut off of .0005 (86).

### Motifs Comparison

In order to compare *de novo* identified motifs, position weight matrices were compared to using TomTom (version 5.5.5). Motifs were compared either against the non-redundant TF database for JASPAR plant TF binding motifs, or compared versus the consensus sequences found in Zenker et al 2024. The most significant motif was used to assign to potential TF families (87–89).

### Data availability

sciATAC-seq data for *Z. mays, S. bicolor, U. fusca,* and *P. miliceum* is found in NCBI under the following bioproject PRJNA1063172. Leaf data for *O.sativa* can be found under the following SRR bioproject PRJNA100757. All scripts used for processing and analyzing data in this manuscript can be found at the following github repository: https://github.com/Jome0169/Mendieta.C4_manuscript. Additionally, all datasets with both MS and BS specific accessibility profiles, their ACRs, as well as their BLASTN relationships can be found on the epigenome browser https://epigenome.genetics.uga.edu/PlantEpigenome. All datasets can be found under the sub-folder Mendieta_et_al.C4_project.

